# Coupling mountain pine beetle and forest population dynamics predicts transient outbreaks that are likely to increase in number with climate change

**DOI:** 10.1101/2023.08.18.553909

**Authors:** Micah Brush, Mark A. Lewis

**Affiliations:** University of Alberta, Department of Mathematical and Statistical Sciences, Edmonton, Alberta, Canada; University of Victoria, Department of Mathematics and Statistics, Victoria, British Columbia, Canada; University of Victoria, Department of Biology, Victoria, British Columbia, Canada

**Keywords:** Mountain pine beetle, *Dendroctonus ponderosae*, Outbreak model, Climate change, Forest structure, Population dynamics

## Abstract

Mountain pine beetle (MPB) in Canada have spread well beyond their historical range. Accurate modelling of the long-term dynamics of MPB is critical for assessing the risk of further expansion and informing management strategies, particularly in the context of climate change and variable forest resilience. Most previous models have focused on capturing a single outbreak without tree replacement. While these models are useful for understanding MPB biology and outbreak dynamics, they cannot accurately model long-term forest dynamics. Past models that incorporate forest growth tend to simplify beetle dynamics. We present a new model that couples forest growth to MPB population dynamics and accurately captures key aspects of MPB biology, including a threshold for the number of beetles needed to overcome tree defenses and beetle aggregation that facilitates mass attacks. These mechanisms lead to a demographic Allee effect, which is known to be important in beetle population dynamics. We show that as forest resilience decreases, a fold bifurcation emerges and there is a stable fixed point with a non-zero MPB population. We derive conditions for the existence of this equilibrium. We then simulate biologically relevant scenarios and show that the beetle population approaches this equilibrium with transient boom and bust cycles with period related to the time of forest recovery. As forest resilience decreases, the Allee threshold also decreases. Thus, if host resilience decreases under climate change, for example under increased stress from drought, then the lower Allee threshold makes transient outbreaks more likely to occur in the future.

**Statements and Declarations:** *Competing interests:* The authors declare no competing interests.

*Data availability statement:* Data sharing is not applicable to this article as no datasets were generated or analysed during the current study. Code to produce the figures is available at github.com/micbru/MPBModel/.

## 1 Introduction

The most recent mountain pine beetle (MPB, *Dendroctonus ponderosae*) outbreak has killed more than 18 million hectares of mainly lodgepole pine (*Pinus contorta*) forests in western Canada, with large economic impacts from the loss of merchantable pine (Corbett et al. 2016; Dhar et al. 2016). Climate change has facilitated MPB incursion into novel habitats, and outbreaks are expected to become more frequent and severe (Logan et al. 2003; Mitton and Ferrenberg 2012; Janes et al. 2014; Bentz et al. 2022). MPB is now threatening further eastward spread into the boreal forest and, if successful, could spread across Canada (Safranyik et al. 2010; Cullingham et al. 2011; Burns et al. 2019). Modelling the long-term dynamics of MPB in the face of global warming and changing forest resilience is central to assessing the risk of further spread and informing management strategies.

There are several key biological elements that must be included to accurately model MPB population dynamics. Beetle development largely follows a one-year development cycle, where adult beetles disperse in the summer to colonize new hosts and then lay eggs inside of the trees. The larvae then hatch and overwinter in the tree before pupating in the spring (Safranyik and Wilson 2006). Small trees do not have thick enough phloem to support egg galleries and beetle development, and so beetles tend to attack only older and larger trees (Amman 1972; Safranyik and Wilson 2006). To overcome tree defenses, including toxic resin, beetles must attack trees in large groups as a mass attack (Raffa and Berryman 1983; Safranyik and Wilson 2006; Goodsman et al. 2016). Beetles achieve this at low population densities by aggregating on local scales, with pheromones signalling to other beetles which tree is under attack (Berryman et al. 1985; Safranyik and Wilson 2006). This process generates an Allee effect, where at very low population densities beetles are unable to attack in large enough numbers to overcome tree defenses. Trees that have been successfully colonized by beetles are generally killed. By the following summer, their needles turn red from a lack of moisture and the trees are called “red tops”. Over the next few years, their needles turn gray and fall off. These trees are called “gray snags”.

To this point, models of MPB outbreaks largely focus on accurately capturing the dynamics of a single outbreak (Heavilin and Powell 2008; Goodsman et al. 2016). The model of Goodsman et al. (2016) produces realistic dynamics by including important aspects of MPB biology, including beetle aggregation and a strong Allee effect. However, models of a single outbreak without forest regrowth cannot predict the long-term effect of different control measures, or tree resilience. In particular, as MPB spreads beyond its historical range there is uncertainty in the resistance of novel host species (Cullingham et al. 2011) as well as evidence that tree resistance to MPB is dependent on historical contact (Cudmore et al. 2010). On the other hand, the model of Duncan et al. (2015), which does include forest age structure, does not include important aspects MPB biology such as an Allee effect or beetle aggregation.

Here, we combine elements of the MPB model of Goodsman et al. (2016) with the forest age structure model in Duncan et al. (2015). In doing so, we create a new model that accurately captures both MPB and forest dynamics and therefore allows us to address long-term questions related to beetle population dynamics, particularly under dynamic forest resilience. In Section 2, we present the model. In Section 3, we study fixed points and derive conditions for a positive fixed point to exist, which biologically can be thought of as the conditions required for an outbreak. In Section 4, we analyze the stability of each fixed point and show that a fold bifurcation gives rise to the non-trivial fixed points. In Section 5 we present numerical simulations of our model and consider the case where MPB immigrates into a stand without an existing MPB population.

## 2 Specifying the model

In this section, we derive the model and define the relevant variables. Because of the annual life cycle of MPB, the model is discrete in time and all variables are indexed with a time *t* that indicates the year. Table 1 provides a summary of the variables and parameters in the model.

**Table 1.** Variables and parameters in the model.

| Variable | Description |
| --- | --- |
| $j_{i,t}$ | Juvenile trees in the $i$th age class in year $t$ |
| $J_t$ | Total juvenile trees in year $t$, $J_t = \sum_{i=1}^N j_i$ |
| $S_t$ | Susceptible trees in year $t$ |
| $B_t$ | Beetles in year $t$ |
| $I_t$ | Infested (green) trees in year $t$ |
| $m_t$ | Mean number of beetles per susceptible tree, $m_t = B_t/S_t$ |
| Parameter | Description |
| $T$ | Total number of trees in the stand |
| $N$ | Number of juvenile age classes |
| $s$ | Juvenile annual survival |
| $c$ | Beetles produced per successfully infested tree |
| $k$ | Beetle aggregation |
| $\varphi$ | Threshold for successful beetle attack |
| $\kappa$ | Beetle aggregation in the Hill approximation |
| $\Phi$ | Threshold for successful beetle attack in the Hill approximation |

### 2.1 Definition of variables

We follow Duncan et al. (2015) in defining the variables we use to track the forest structure and beetle populations. We divide trees into juvenile, susceptible, and infested classes. We further divide juvenile trees into *N* age classes, with each class representing a year of growth. We let *j*_*i,t*_ be the number of trees in the *i*th juvenile age class in the spring of year *t* (the number of juvenile trees is measured in terms of the shade equivalent of adult trees, as we assume the forest is light limited). The total number of juvenile trees in the spring is then 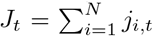 We let *S*_*t*_ be the number of trees large enough to be susceptible to MPB infestation in the spring of year *t*. We let *B*_*t*_ be the number of beetles that emerge in the summer of year *t* from last year’s infested trees. We let the number of newly infested trees in the summer of year *t* be *I*_*t*_.

Because pine forests are shade intolerant (Safranyik and Wilson 2006; Duncan et al. 2015), we track infested trees until they no longer shade the forest floor. We assume that trees that have been infested more than two years ago do not have enough needles to produce substantial shade (Safranyik and Wilson 2006). The number of red tops in the spring of year *t* is then given by *I*_*t−*1_, or the number of infested trees from the previous summer, and the number of gray snags in the spring of year *t* is given by *I*_*t−*2_.

### 2.2 Forest dynamics

The forest dynamics are modeled as in Duncan et al. (2015) (see Figs. 1 and 2 there for life cycle diagrams). We assume that juvenile trees have some natural probability of survival *s* from one year to the next, with their mortality then equal to *d* = 1 − *s*. If they survive, they advance to the next age class. This means that

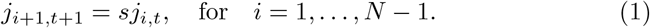

Juvenile trees that survive from the *N* th age class become susceptible trees. We assume that susceptible trees have no natural mortality, and die only upon being infested by beetles. Thus, in any year, susceptible trees either escape infestation and remain susceptible, or are infested by beetles. The number of susceptible trees in the following year is then given by

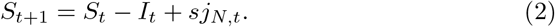

Note that the number of infested trees must be less than the number of susceptible trees in a year, and so this equation remains positive. Mathematically, the equation we derive in the following section for *I*_*t*_ guarantees that *I*_*t*_ ≤ *S*_*t*_.

**Fig. 1.**
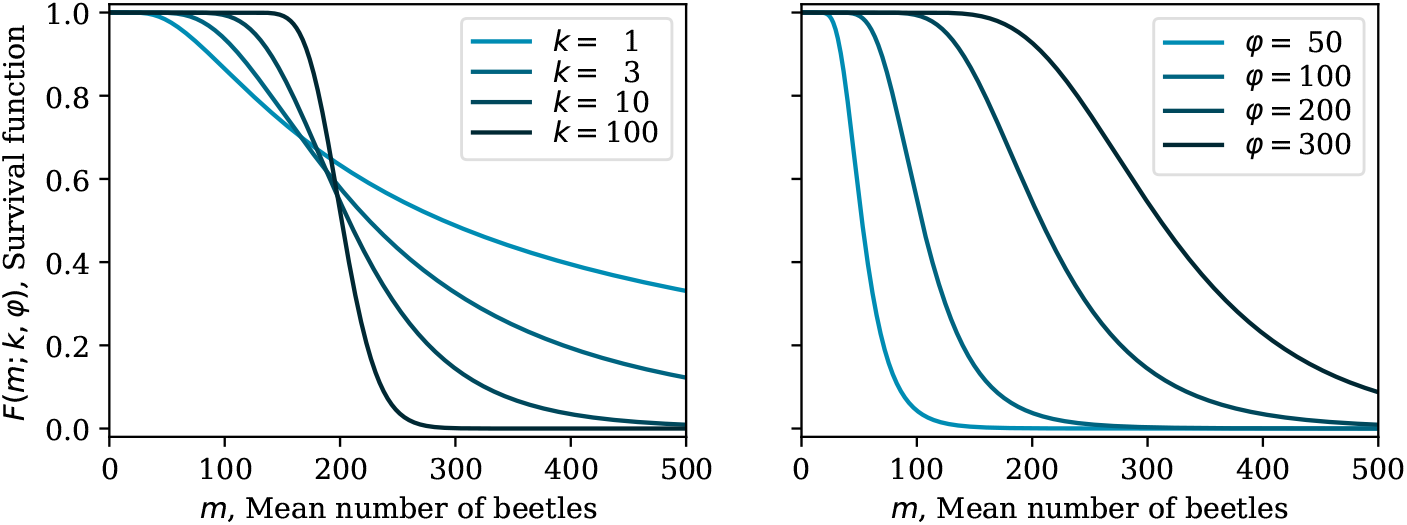
The survival function *F* (*m*_*t*_; *k*, φ) for different values of the aggregation parameter *k* and the threshold *φ*

**Fig. 2.**
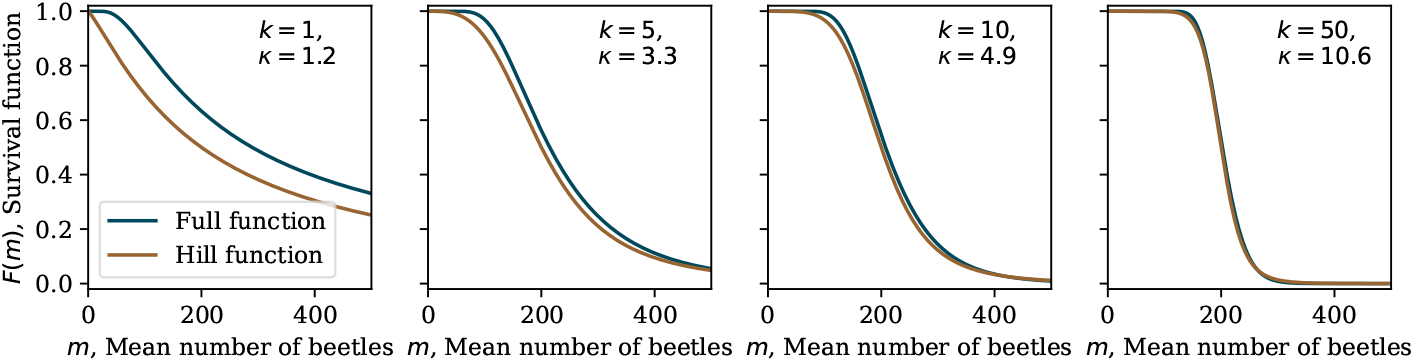
Comparison of the full function 1 − *F* (*m*; *k*, φ) to the Hill function approximation. We set Φ = φ = 200 and set *κ* by minimizing the difference between the functions with *k* = 1, 5, 10, 50

For seedling recruitment, we assume that seedlings germinate wherever there is available light, and that growth is not limited by seeds (as in Duncan et al. (2015) and Křivan et al. (2016)). These are reasonable assumptions as MPB outbreaks are known to provide openings for new growth (Amman 1977; Axelson et al. 2009), and for lodgepole pine in particular, the dominant MPB host, there is typically a very large seed bank as cones are serotinous and do not germinate until after disturbance. These disturbances are typically fire (Johnson and Fryer 1989), though MPB attacks may also trigger seed release (Teste et al. 2011). We therefore allow new seedlings to take the place of all juvenile trees that died in the previous year as well as all gray snags old enough to no longer shade the forest floor. The number of new seedlings is then given by

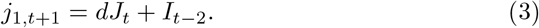

Because of our assumption that seedlings replace all trees that no longer shade the forest floor, the total number of trees *T* inventoried in the spring is conserved from year to year. The total number of trees is given by

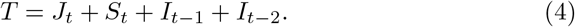

To see that this is conserved, set *T* = *J*_*t*+1_ + *S*_*t*+1_ + *I*_*t*_ + *I*_*t−*1_ and substitute (1), (2), and (3).

### 2.3 Beetle dynamics

Our model for the number of emerging beetles is based on Goodsman et al. (2016), but here we make use of the general site based framework following Brännström and Sumpter (2005) and Anazawa (2009). We can use this frame-work to derive models mechanistically from first principles, which gives them biological grounding. Consider a large number of sites *S*_*t*_ (susceptible trees) at time *t*. Let *p*(*i*; *B*_*t*_) be the probability of *i* individuals at a site given *B*_*t*_ total individuals (beetles). We assume that all sites are equally suitable. Let *n*(*i*) be the number of individuals that emerge from infested sites the following year. Then, the total number of individuals that emerge in the following year, *B*_*t*+1_, from the *S*_*t*_ sites in the previous year is given by

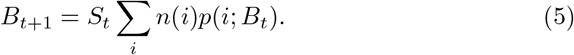

This equation generally leads to different functional responses depending on what we assume for the distribution of individuals over sites and the number of individuals emerging from each site (Brännström and Sumpter 2005; Anazawa 2009).

We can derive the functional form for infestation in Goodsman et al. (2016) in this framework by assuming that the beetles are aggregated and that there is some threshold at each site below which the beetles cannot successfully reproduce. This models the beetle aggregation due to pheromones as well as the natural defenses of the tree, which are key biological aspects of beetle population dynamics. Note that we track only female beetles, as they infest the trees.

To model aggregation, we assume (as in Goodsman et al. (2016)) that the beetles are distributed according to a negative binomial distribution with mean *m*_*t*_ = *B*_*t*_*/S*_*t*_ and aggregation parameter *k*, where low *k* corresponds to high aggregation and high *k* corresponds to low aggregation,

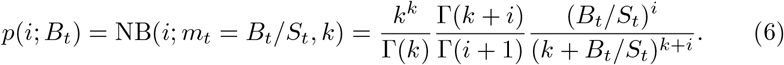

We then assume that the beetle productivity *c* is constant as long as the number of beetles attacking a tree is above the threshold φ required to overcome tree defenses. This is equivalent to assuming contest competition, where by contrast scramble competition would lead to overcompensation at high beetle densities (Brännström and Sumpter 2005; Anazawa 2009). Then,

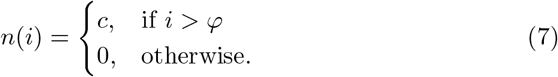

Thus, each tree produces either *c* or 0 beetles, and the number of trees killed depends on how the beetles distribute themselves amongst the trees according to *p*(*i*; *B*_*t*_).

Inserting the biologically motivated (6) and (7) in (5), we find

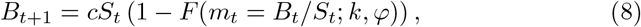

where

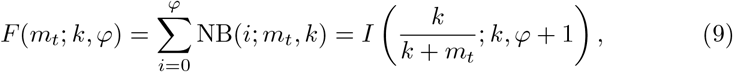

and *I*(*x*; *a, b*) is the cumulative distribution function of the negative binomial distribution, which is the regularized incomplete beta function. Note that here we have changed the order of the variables and parameters in the function *F* compared to the negative binomial distribution. This is convenient as in the model we take the aggregation *k* and the threshold φ to be fixed parameters, while the number of beetles is the dependent variable and changes over time with the population dynamics of *B* and *S*.

As in Goodsman et al. (2016), we identify this function *F* as a survival function for the susceptible trees. That is, the number of surviving trees in each year is given by *F* (*m*_*t*_)*S*_*t*_. We plot the function *F* for several values of *k* and φ in Fig. 1 to show how different levels of aggregation and different thresholds for tree defenses change the fraction of trees that survive. Since the number of surviving trees is given by *F* (*m*_*t*_)*S*_*t*_, the number of infested trees *I*_*t*_ can be identified as

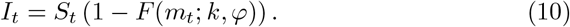

nd so, substituting into (8), we can write

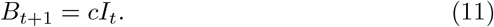

### 2.4 Approximation of survival function

The survival function derived in the previous section can be quite challenging to work with analytically. To make progress analyzing this model, it is useful to approximate the survival function with a Hill function, as in Křivan et al. (2016). We denote this approximation as 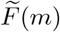 which is equal to

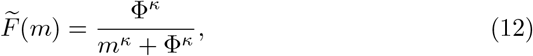

or alternatively

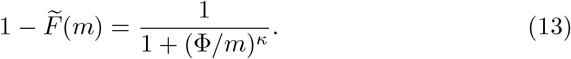

The parameter Φ approximates the threshold number of beetles φ, and *κ* approximates the beetle aggregation parameter *k*, with lower *κ* corresponding to higher levels of aggregation and with beetles spread uniformly in the limit as *κ* → ∞. These functions do not correspond perfectly, but the approximation works well for many values of the parameters by setting Φ = *φ* and then finding *κ* by minimizing the square of the difference for fixed *k*, ie. Minimizing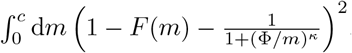 We compare the Hill function and the full survival function for a few different values of *k* in Fig. 2.

Note that this approximation does not work well for high aggregation (small *k*), as in this limit the Allee threshold becomes less important. When fitting their model to infestation data from Willmore Wilderness Park in Alberta, Canada, Goodsman et al. (2016) find a best fit value of *k* ≈ 0.003. We do not find that this is a good direct comparison to the model presented here as this is for a spatial extension of their model with beetle dispersal and immigration, and does not allow for new trees to enter the susceptible class (ie. *s* = 0). We note additionally that fitting the aggregation parameter is sensitive to the spatial grain and scale of the data, and the best fit value may prefer larger values at different spatial scales. In what follows we make use of the Hill function approximation and assume that beetles are not strongly aggregated. However, similar analysis could be carried out with a different approximation in the case *k* ≪ 1 (Appendix A).

### 2.5 Model summary

To finalize the equations defining the model, we first obtain an expression for *S*_*t*+1_ that does not depend on *I*_*t*_ by substituting (10) into (2),

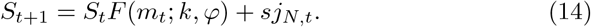

We then rewrite (3) so that the right-hand side depends only on quantities observed at time *t*. We use the conservation of the total number of trees *T* (4) and substitute for *I*_*t−*2_. We then use *B*_*t*_*/c* = *I*_*t−*1_ to substitute for *I*_*t−*1_ andobtain

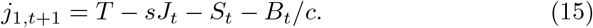

In summary, our model consists of (1), (8), (14), and (15), which we write again here for clarity as

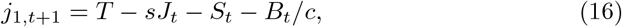

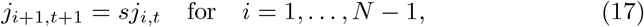

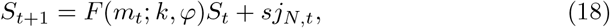

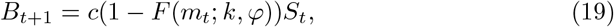

Where 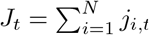

### 2.6 Nondimensionalization

We can now nondimensionalize the model. Let

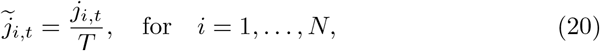

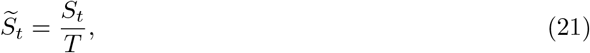

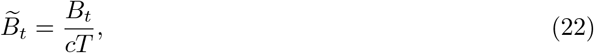

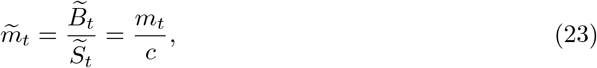

and rewrite the equations above after dividing through by *T* to obtain

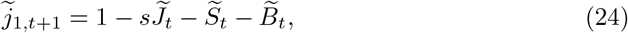

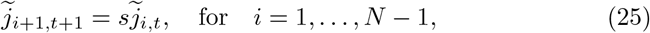

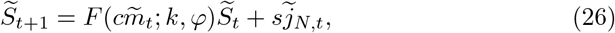

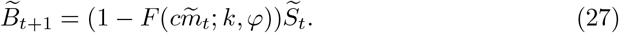

This removes *T* as a parameter and thus 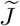 and 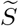 represent the fraction of total trees (4) rather than the number of trees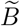 has also been normalized by the number of beetles that emerge at a site, *c*, so it can now be thought of as the fraction of infested trees from the previous year by using (11).

To further reduce the number of parameters, we would have to rescale either φ or *k* with *c* to remove one as a parameter. This is not possible with the definition of *F* in (9). However, with the Hill approximation outlined above in (13), we find that Φ*/c* appears together and can therefore be defined as a new parameter, *θ* = Φ*/c*, which is a measure of host resilience. The nondimensionalized equations are then

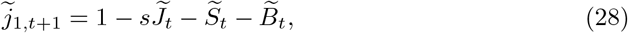

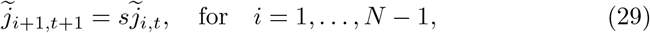

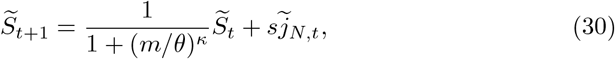

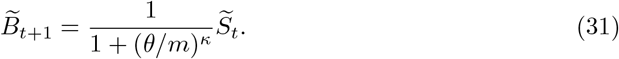

In the analysis that follows, we use the equations that carry dimension (16)– (19) as well as the nondimensionalized equations (28)–(31) because the full survival function *F* cannot be nondimensionalized in this way. Nonetheless, we drop the tildes for notational simplicity and specify when we make use of the nondimensionalized equations.

This nondimensionalization is particularly useful biologically because both the threshold Φ and the number of emerging beetles *c* could be modified under changes in tree resilience, but at this point it is not well understood how these vary in trees with genetic resilience (Six et al. 2018). This nondimensionalization shows that either decreasing the threshold or increasing the number of emerging beetles have the same effect on the dynamics.

## 3 Fixed points

In this section, we solve for the fixed points of the model. We show that there is always a trivial fixed point without beetles and that there can be one or two additional non-trivial fixed points. We then solve for the values of the variables at these non-trivial fixed points. Finally, we use the Hill function approximation to derive the condition for the appearance of the non-trivial fixed points and at what critical value they appear.

### 3.1 Derivation of the fixed points

We begin by dropping the time indices on (16)–(19). When there are no beetles, *B* = *m* = 0 and *F* (0) = 1, which inserting into (2) gives *j*_*N*_ = 0 and thus *j*_*i*_ = *J* = 0, and finally using (4) gives *S* = *T* . Thus, at the trivial fixed point there are no beetles and all trees are susceptible. Biologically, this is because the model does not have mortality from causes other than pine beetle, and so in the absence of beetles the trees all reach their susceptible age. In reality, lodgepole pine are an early successional species often replaced by more shade tolerant trees, and their dynamics are highly fire dependent (Johnson and Fryer 1989; Axelson et al. 2010; Taylor and Carroll 2003). However, this fixed point makes some sense in a stand of pure lodgepole pine with very slow succession and in the absence of fire.

More generally, at all fixed points (19) becomes *B* = *c*(1 − *F* (*m* = *B/S*))*S*, and dividing by *S* gives the mean number of beetles per tree at a fixed point,

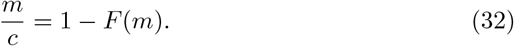

This equation is important for understanding the dynamics of the model. In general it must be solved numerically, but we can show that it has either 1, 2, or 3 roots, including the trivial root at *m* = 0. To understand this graphically, we plot the function 1 − *F* (*m*) and *m/c* together in Fig. 3 for different parameter values showing the number of roots. Intuitively, the function 1 − *F* (*m*) has a monotonically increasing sigmoidal shape as it is 1 minus the cumulative distribution function of the negative binomial distribution (see Fig. 1), and thus a straight line that goes through 0 intersects this curve at the origin, and then either never again, once, or twice more depending on the parameter values. Mathematically, note that *F* ^*′*^(0) = 0, and so the line *m/c* is above 1 − *F* (*m*) for small *m*, and because of the sigmoidal shape of 1 − *F* (*m*) the lines intersect once or twice more. More formally, we can show that the second derivative of 1 − *F* (*m*) − *m/c* has a unique zero for *m >* 0, and so 1 − *F* (*m*) − *m/c* = 0 has at most three roots for *m >* 0. Note that in the case of the Hill function approximation, we can show this in a different way by deriving the conditions for the appearance of additional roots (see Section 3.2).

**Fig. 3.**
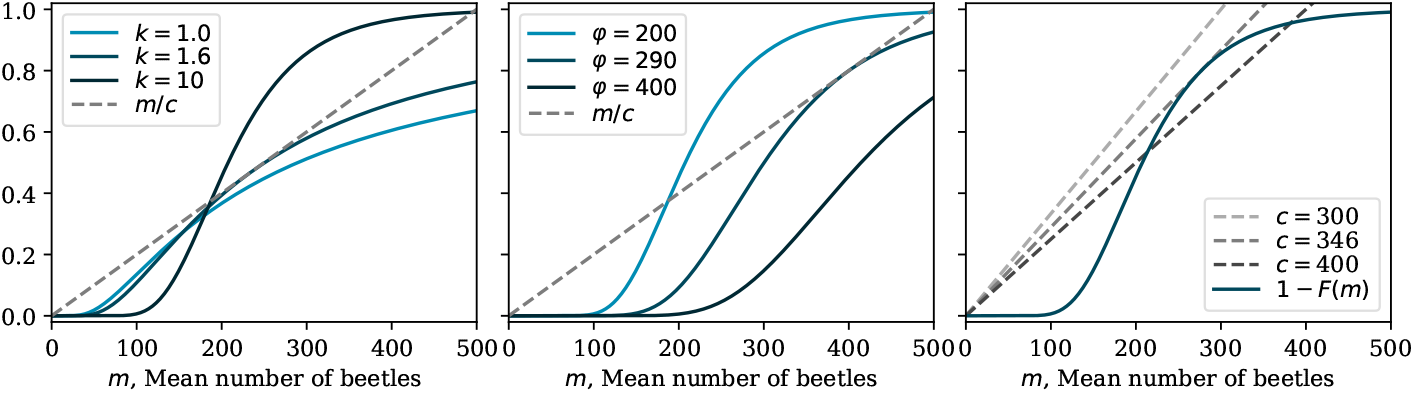
Graphical solutions for the number of roots with changing model parameters *k*, φ, and *c*. The dashed lines show *m/c* and the solid lines show 1 − *F* (*m*; *k, φ*), where the parameter being varied is shown in the legend. When not varied, *k* = 10, *φ* = 200, and *c* = 500. The intermediate curve in each case shows the value of the parameter where there is exactly one non-trivial root, the darkest curve shows an example where there are two non-trivial roots, and the lightest curve shows an example where there are no non-trivial roots

To solve for the juvenile age classes at a fixed point, first note that the total number of trees (4) at a fixed point is *T* = *J* + *S* + 2*I* = *J* + *S* + 2*B/c*, and so we can write (16) for *j*_1_ as

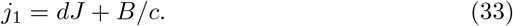

We can then solve for the total number of juvenile trees *J* at a fixed point. From (17), *j*_*i*_ = *s*^*i−*1^*j*_1_, and so

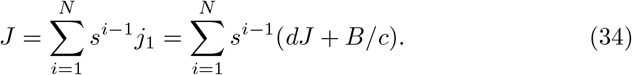

Solving for *J*, we find

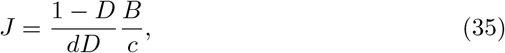

where *D* = *s*^*N*^ . We substitute this back into our expressions for *j*_1_ to find *j*_1_ = *B/*(*cD*) and derive a formula for the number of juveniles in age class *i*. Having specified *j*_*i*_ and *m* at the fixed points, we can use the equation for *T* (4) to solve uniquely for *S* and *B* after solving the equation for *m*. The final set of equations to obtain the fixed points of the model are

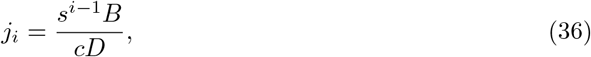

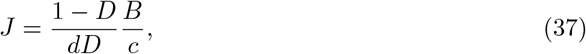

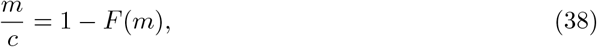

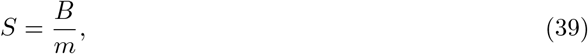

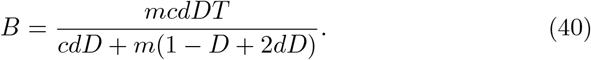

### 3.2 Conditions for non-trivial fixed points using the Hill function approximation

Given that *F* is a complicated function, the equation for *m* at the fixed point (32) cannot typically be solved analytically. However, we can use the Hill function approximation introduced earlier to approximate the parameter values required for non-trivial fixed points. The existence of these non-trivial fixed points is required for pine beetle outbreaks to occur as without them the population goes towards the trivial fixed point and the beetles die out (see Section 4).

Using the Hill function approximation (13), the equation for the fixed points (32) becomes

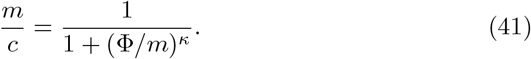

Assuming m ≠ 0 (ie. assuming beetles are present), this can be rewritten as *m*^*κ*^ − *cm*^*κ−*1^ + Φ^*κ*^ = 0. When this equation has roots with *m >* 0, there exist non-trivial fixed points with a positive number of beetles. Therefore we want to find the conditions on this equation such that there are roots where *m >* 0. Let

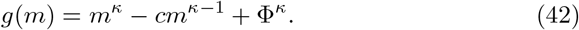

The derivative of *g*(*m*) is then

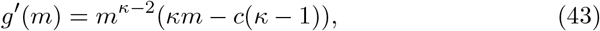

which has a unique zero at

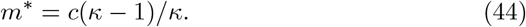

Since *g*(0) *>* 0, and lim_*m→∞*_ *g*(*m*) = lim_*m→∞*_ *m*^*κ*^ *>* 0, and the derivative of the function has a unique zero at *m*^*∗*^, the function only has roots if *g*(*m*^*∗*^) ≤ 0. More precisely, *g*(*m*) has one root if *g*(*m*^*∗*^) = 0, two roots if *g*(*m*^*∗*^) *<* 0, and no roots if *g*(*m*^*∗*^) *>* 0. Substituting *m*^*∗*^ in to *g*(*m*), we find

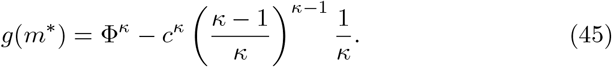

Rearranging (45) with *g*(*m*^*∗*^) *<* 0 then gives the condition on the parameters such that we have two roots,

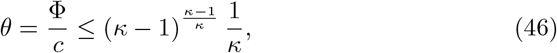

where *θ* = Φ*/c* is the nondimensionalized resilience parameter introduced earlier. When this condition is satisfied, there are positive, non-trivial solutions to (41), and therefore fixed points with a positive population of beetles. This means that it possible for beetle outbreaks to occur. We have exactly one root when this condition is equal to zero. This occurs at

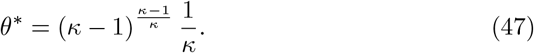

Biologically, the condition in (46) makes sense in the limit of large *κ* (low aggregation) as the condition becomes Φ *< c*. In other words, the number of beetles that emerge per tree must be greater than the threshold when the beetles are uniformly distributed. In the limit of small *κ* (strong aggregation), the behaviour is initially somewhat counter intuitive. Because the right-hand side of (46) increases monotonically from 0.5 to 1 for *κ >* 2, as beetles become more aggregated the right-hand side decreases. This puts a stricter upper bound on the ratio of the threshold and beetle productivity as the beetles become more aggregated. We can see this in Fig. 3 for the full function with changing *k*, where as the beetle aggregation increases (decreasing *k*) with other parameters fixed, the non-trivial fixed points disappear. This behaviour can be understood because the number of beetles produced per infested tree is constant regardless of how strongly the beetles aggregate on a single tree. This means that as beetles aggregate more strongly onto fewer trees, fewer beetles are produced the following year.

## 4 Stability analysis

In this section, we analyze the stability of the fixed points found in the previous section. We show that the trivial fixed point is stable with both the full function and the Hill function approximation. Using the Hill function approximation, we then show that the smaller (in terms of number of beetles) of the two non-trivial fixed points is unstable. Finally, again using the Hill function approximation, we show that a fold bifurcation occurs at *m*^*∗*^ and *θ*^*∗*^, and therefore that in the local neighbourhood of the critical point the upper of the two non-trivial fixed points is stable.

To analyze the local stability of the model, we expand (16)–(19) about the fixed points. Writing the variables as a vector *x* = (*j*_1_, *j*_2_, …, *S, B*), we rewrite the model as *x*_*t*+1_ = φ (*x*_*t*_), where the function φ is determined by the right-hand side of (16)–(19). Expanding around a fixed point *x* = *x*_0_ gives

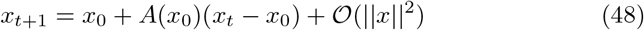

where *A* is the Jacobian matrix given by the expansion of φ at the fixed point. We can write the Jacobian for the model explicitly as the *N* + 2 dimensional square matrix

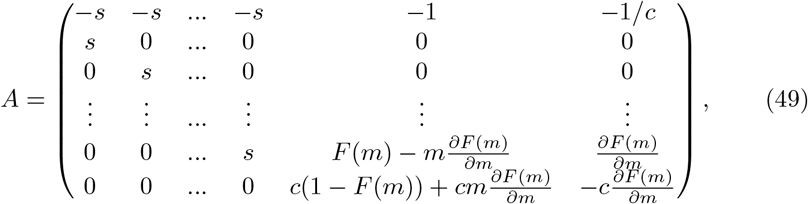

where the mean number of beetles at the fixed point is *m*.

We can write the characteristic polynomial *P* of this matrix analytically, where

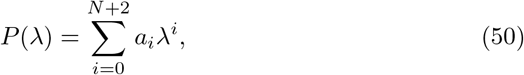

and the *a*_*i*_ coefficients are

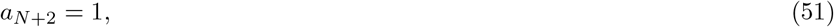

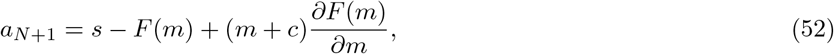

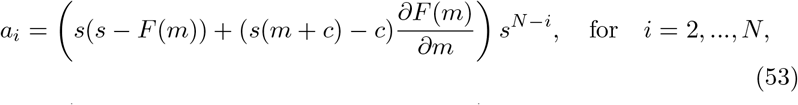

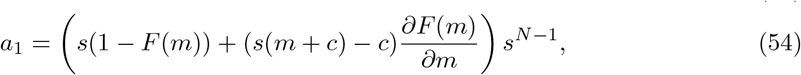

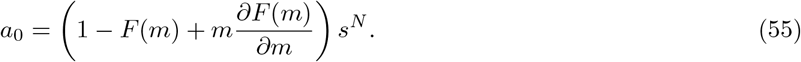

### 4.1 Stability of the trivial fixed point

We here use Rouché’s theorem to show that the trivial fixed point is always stable for both the full function and the Hill function approximation. Biologically, this means that when the forest consists of susceptible trees and no beetles it is stable.

Rouché’s theorem states that for two complex functions *g*_1_ and *g*_2_ in the region *K* with closed contour *∂K*, if |*g*_2_| *<* |*g*_1_| on the contour *∂K*, then *g*_1_ and *g*_1_ + *g*_2_ have the same number of zeroes inside *K*. This theorem can be used to bound roots of polynomials. To bound the roots of the characteristic polynomial *P* (50), we write 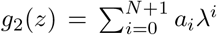 and 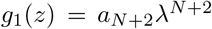 so *P* = *g*_1_ + *g*_2_. We know that *g*_1_ has *N* + 2 roots at *λ* = 0. If |*g*_2_| *<* |*g*_1_| on the boundary of the disk centered on the origin with radius *R* (*R >* 0), then by Rouché’s theorem *P* = *g*_1_ + *g*_2_ also has *N* + 2 roots inside the disk. Writing |*g*_2_| *<* |*g*_1_|, we find

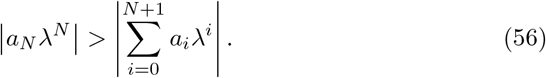

On the boundary 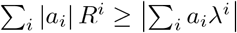 since *R* is the maximum value of *λ* on the disk, and (56) then becomes

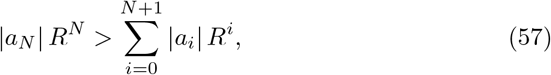

which is therefore a sufficient condition for the roots of *P* to be bounded within the disk with radius *R*.

We can apply this approach to the analysis of the stability of a fixed point. If all roots of the characteristic equation are less than 1, the fixed point is locally stable. This corresponds to the spectral radius of the Jacobian being less than 1. Thus, if *P* satisfies (57) with *R* = 1 at a fixed point, it has has all *N* + 2 roots within a disk of radius 1, and therefore that fixed point is locally stable. If the condition in (57) is not satisfied, we cannot determine the stability of the fixed point. We now write the condition (57) with *R* = 1 using the definitions of the *a*_*i*_ for (50) and find

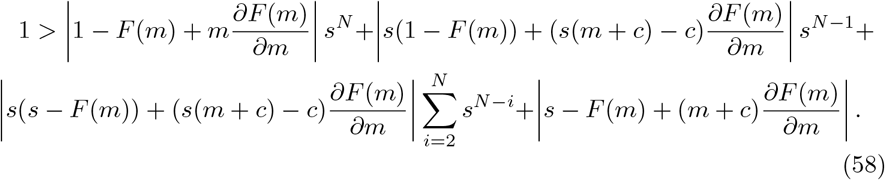

At the trivial fixed point, *m* = 0, *F* (0) = 0, and 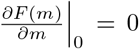 We can evaluate the remaining sum over *s* analytically as 0 *< s <* 1 to simplify (58) and obtain

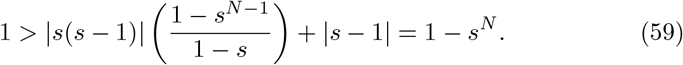

This condition is true given that 0 *< s <* 1, since *s* is the probability of survival of a juvenile tree. Therefore, (57) is satisfied for the trivial fixed point with *R* = 1 and thus by Rouché’s theorem all roots of the characteristic polynomial at the trivial fixed point are within the unit disk. Thus, the trivial fixed point is stable. Biologically, this makes sense as we know the beetles have an Allee effect.

### 4.2 Stability of the non-trivial fixed points

We now turn to the analysis of the non-trivial fixed points. To make progress analytically, we use the Hill function approximation. We show that the smaller (in terms of number of beetles) of the two non-trivial fixed points is always unstable using the Jury stability criterion.

In *n* dimensions, the Jury stability criterion requires the construction of a table with 2*n* − 3 rows to test *n* + 1 conditions on the characteristic polynomial *P* at a fixed point. If all of the conditions are satisfied, the fixed point is stable. This becomes difficult analytically with higher dimensional systems. However, as soon as any condition is not met, we can conclude the fixed point is unstable. In this case, we use only the first condition, which requires that *P* (1) *>* 0. For the characteristic polynomial in (50), we find

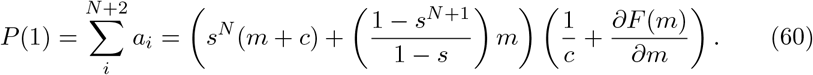

The term inside the first parentheses in (60) is always positive, as 0 *< s <* 1, *m* ≥ 0, and *c >* 0. Therefore to determine the sign of (60), we only need to consider the term inside the second parentheses. The requirement that *P* (1) *>* 0 can therefore be written as

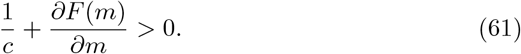

To make further progress, we now use the Hill function approximation for *F*, 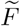 (12). The partial derivative of the Hill function approximation is

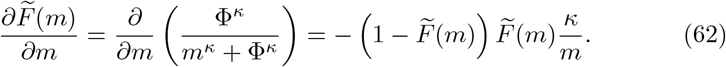

At a fixed point, we have 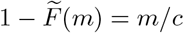 and so we can rewrite (62) as

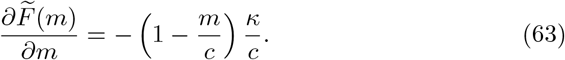

We substitute (63) in to (61) to rewrite the condition in terms of *m* as

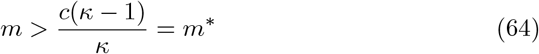

where *m*^*∗*^ is given by (44) and is the value where we have exactly one non-trivial fixed point.

Thus, the condition that *P* (1) *>* 0 is only satisfied if *m > m*^*∗*^. We have two non-trivial fixed points, one with *m < m*^*∗*^ and one with *m > m*^*∗*^. Therefore, the smaller of these two fixed points is always unstable. Note that we cannot conclude anything about the stability of the larger fixed point from this case without testing other conditions.

### 4.3 Bifurcation analysis

We here show that our model undergoes a fold bifurcation at *m* = *m*^*∗*^ by making use of center manifold theory. As in the examples in the previous paragraph, it is the appearance of these new fixed points that give rise to outbreaks in our model. This bifurcation results in one stable and one unstable fixed point. As we have already shown that the smaller of the two fixed points is unstable, we conclude that the larger fixed point is stable in the immediate neighbourhood of the bifurcation.

To setup the bifurcation analysis, we start with the nondimensionalized equations using the Hill approximation (28)–(31). We define shifted variables, which we denote with a bar, such that they are 0 at the critical fixed point when *m* = *m*^*∗*^ (44) with *θ* = *θ*^*∗*^ (47). The shifted variables are

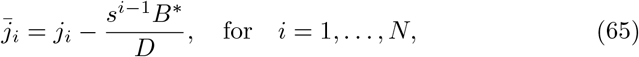

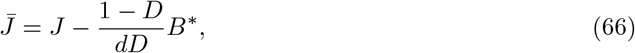

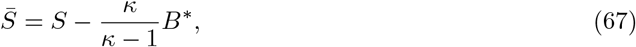

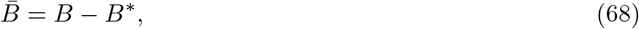

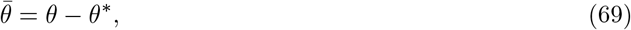

where *B*^*∗*^ can be obtained by setting *m* = *m*^*∗*^ in (40). Note that when *m* = *m*^*∗*^ and *θ*=*θ*^*^,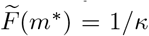 Substituting these shifted variables in to our model (28)–(31) gives the shifted equations

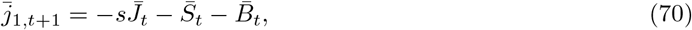

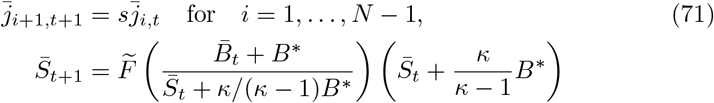

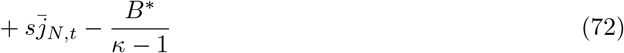

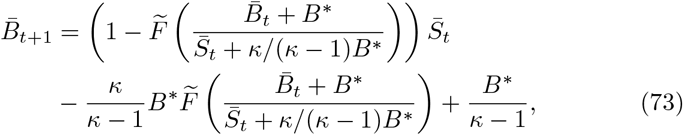

where 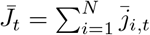 We then write the variables as a vector 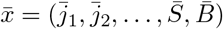 and rewrite the shifted equations as 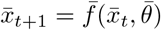 . We Taylor expand 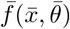 around the critical fixed point 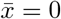 with 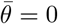 to obtain

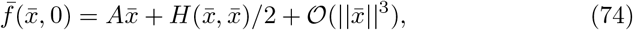

where *A* is the Jacobian evaluated at 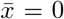 and the *i*th element of *H*(*x, y*) is defined as

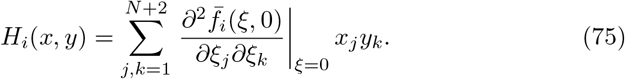

With the notation established, we now make use of center manifold theory following Kuznetsov (2004) to show that the model undergoes a fold bifurcation at the critical fixed point 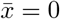 with 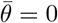

First, in order to show that the dynamics of the system can be reduced to a 1-dimensional center manifold, we must show that the Jacobian at the critical point has a simple eigenvalue of *λ* = 1. We calculate the Jacobian at the critical point and find

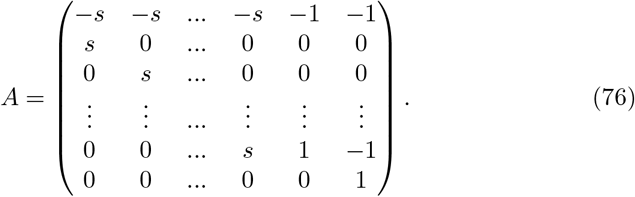

The characteristic equation is then

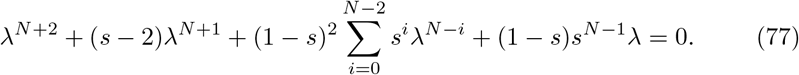

Taking out the factors corresponding to the roots at *λ* = 0 and *λ* = 1 leaves

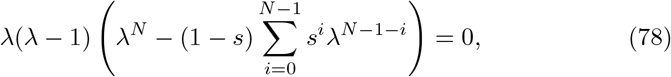

and so the Jacobian has a root at *λ* = 1. We must then show that this root is a simple root, which we do by showing the polynomial 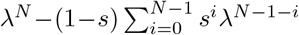 in (78) does not have *λ* = 1 as a root. We apply Rouché’s Theorem as in (57) to show that all of the roots are less than one as long as

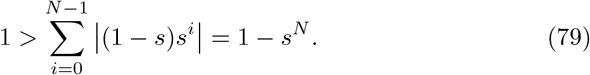

This conditions is satisfied as 0 *< s <* 1, and so all of the remaining roots of (78) are less than one. Therefore, we have shown that the Jacobian has a simple root of *λ* = 1 at the critical point, and thus that the dynamics can be reduced to the 1-dimensional eigenspace spanned by the eigenvector associated with *λ* = 1. Next, we have to show that the dynamics of this reduced system have a fold bifurcation at the critical point. We define *p* and *q* as the adjoint eigenvector and the eigenvector corresponding to *λ* = 1, respectively, (ie. *A*^*T*^ *p* = *p* and *Aq* = *q*), and normalize them such that ⟨*p, q*⟩ = 1, where ⟨·, ·⟩ is the scalar product. We can then show the restriction to the center manifold takes the form 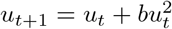, where *u*_*t*_ lives on the 1-dimensional center manifold and *b* can be calculated as

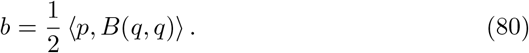

As long as b ≠ 0, the system undergoes a fold bifurcation at the critical point.

The eigenvector corresponding to *λ* = 1 is

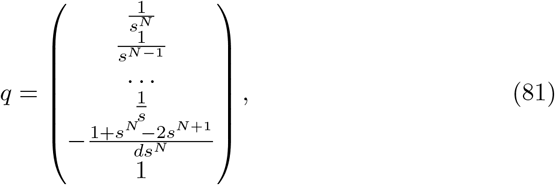

and the normalized adjoint eigenvector corresponding to *λ* = 1 is

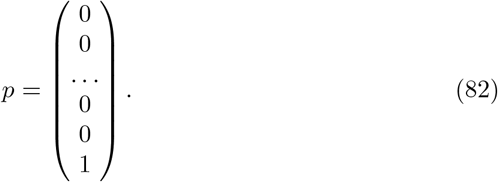

Given that the only nonzero element of *p* is the final element, we only need to calculate the final element of *B*(*q, q*) in order to calculate the coefficient *b* in (80). The final element of *B*(*q, q*) is

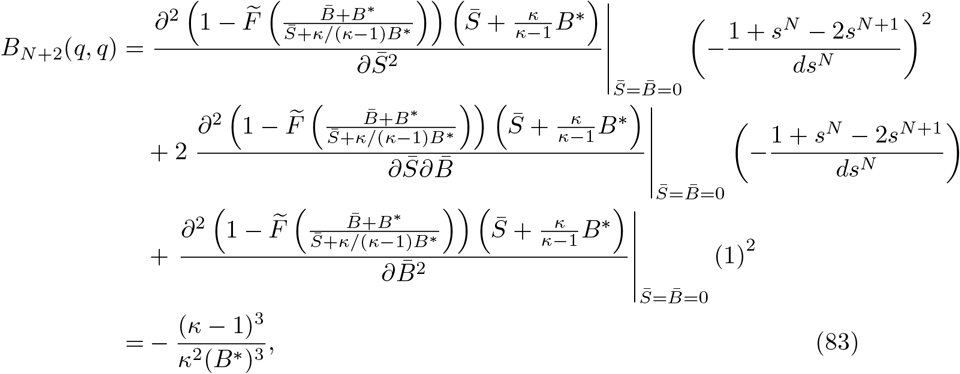

and so

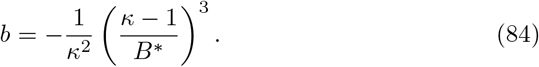

This is not zero as long as *κ >* 1 and we have a non-zero number of beetles, both of which are true at the non-trivial fixed point. Therefore, the system undergoes a fold bifurcation at the critical point.

To summarize, we have shown that at the critical point *m* = *m*^*∗*^ with *θ* = *θ*^*∗*^, the Jacobian has a simple eigenvalue *λ* = 1 and therefore the dynamics can be reduced to a one dimensional center manifold with 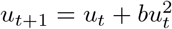 We have also shown that the coefficient *b* is not zero, and so there is a fold bifurcation at the critical point. With *θ* as the bifurcation parameter, this means we gofrom zero positive fixed points for *θ > θ*^*∗*^ to two positive fixed points for *θ < θ*^*∗*^, where one is stable and one is unstable near the critical point. Because we have shown that the smaller of these two fixed points is always unstable, the larger fixed point that appears is stable in the neighbourhood of the critical point.

## 5 Numerical results

In this section, we analyze the dynamics and stability of the model numerically. We test the stability of the non-trivial fixed points with the original *F* function and compare it to our analytical results using the Hill function approximation. We then simulate the system with different initial conditions to investigate its transient dynamics as it approaches the stable fixed points.

In all of the following simulations and figures, we fix the number of years before trees become susceptible at *N* = 50 and the juvenile survival rate at *s* = 0.99, as in Duncan et al. (2015). The biological plausibility of the number of years before trees become susceptible can be seen in Fig. 8 of Safranyik and Wilson (2006), where they find a few attacked trees below age 50 but many more between 50 and 60. The plausibility of the juvenile survival rate can be seen in Fig. 3 in Johnson and Fryer (1989), where the lodgepole pine mortality per 5 year interval for stands less than 50 years old varies between 0.001 and 0.1. Alternatively, we can calculate *s* ≈ 0.99 using 15 year seedling survival rates for lodgepole, hybrid, and jack pine from Rweyongeza et al. (2007). The qualitative results discussed below are not sensitive to these exact values.

### 5.1 Numerical stability of the non-trivial fixed points

We first study the bifurcation in *θ* numerically and compare the results obtained with the Hill function approximation 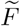 (12) and the original *F* function (9). To make this comparison, we use dimensional parameters as the nondimensionalization we used relies on the Hill function approximation (see Section 2.6). We fix *c* = 500, which is similar to the value derived from data in Goodsman et al. (2016), and *k* = 10, which gives *κ* ≈ 4.9 when minimizing the difference between the functions (see Section 2.4). We then vary the threshold parameter (φ in the original function and Φ for the Hill function approximation) in integer increments from 1 to *c*. We use integer increments as the number of attacking beetles must be integer valued (mathematically this threshold must be integer valued given the definition as the sum over the negative binomial distribution in (9)). Figure 4 shows the resulting bifurcation diagram, where solid lines are stable and dashed lines are unstable. The stability is determined numerically by calculating the spectral radius of the Jacobian. The bottom curve here corresponds to the trivial fixed point, and the additional curves that appear below a critical threshold correspond to the non-trivial fixed points. The non-trivial fixed points were calculated by numerically solving (32) with both the original function and the Hill function. We note that the original function and the Hill function have similar stability properties, even if the bifurcation occurs at a slightly lower value of the threshold parameter for the original function.

**Fig. 4.**
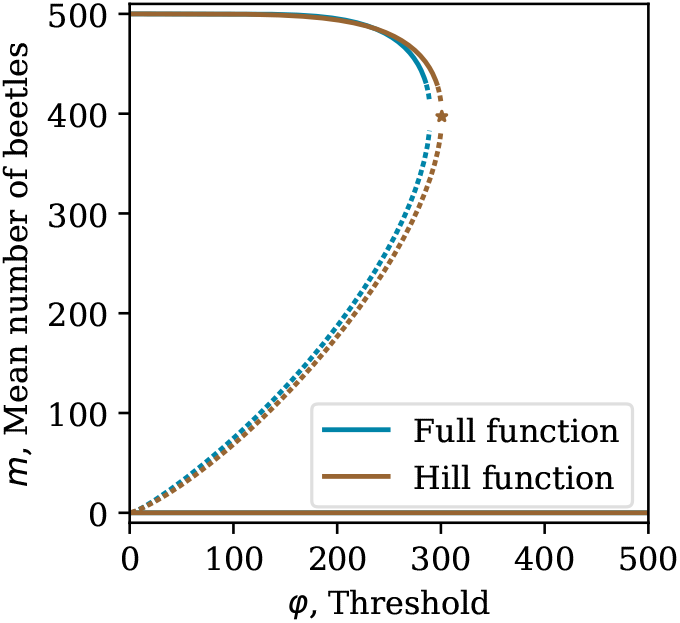
The bifurcation diagram in the threshold parameter with the numerical results of the original survival function *F* (*m*) and the Hill function approximation. We set *k* = 10 and *c* = 500 for the full function, and for the Hill function we set φ = Φ and minimize the difference between the functions to obtain *κ* ≈ 4.9. Solid lines are stable and dashed lines are unstable. The star indicates the critical point for the Hill function approximation with *m*^*∗*^ = *c*(*κ* − 1)*/κ* and *θ*^*∗*^ = Φ*/c* = (*κ* − 1)^(*κ−*1)*/κ*^*/κ*

We next compare the numerical results to our analytic results. First, we notice that the trivial fixed point is indeed stable for all parameter values for both the original function and the Hill function approximation. In the case of the Hill function approximation, we find that the derived critical values of *m*^*∗*^ (44) and *θ*^*∗*^ (47) (marked by a star in Fig. 4) accurately predict the value of the threshold where the non-trivial fixed points appear. Additionally, the smaller of the two-non trivial fixed points appears unstable for all parameter values where it exists, as predicted.

However, the numerical results seem to indicate that the larger of the nontrivial fixed points is in fact unstable near the critical point in both cases, while the analytical results show that this point should be stable, at least near the critical point when using the Hill function approximation. We plot the spectral radius of all three fixed points in the Hill function case very near to the critical point in Fig. 5. Note that here we have nondimensionalized so the resilience parameter *θ* is now continuous, and have fixed the aggregation at *κ* = 5. We see that in the immediate neighbourhood of the critical point, when the non-trivial fixed points first appear, the upper fixed point is indeed stable for a very small range of the bifurcation parameter *θ*. The spectral radius then increases slightly above one for a small range of *θ* before returning below one as *θ* decreases. We note that the real and imaginary parts of the dominant eigenvalue are both positive over the range of *θ* in Fig. 5. Importantly, The parameter range where the spectral radius is greater than one is small, here extending only from *θ* ≈ 0.598 − 0.606. With *c* = 500, this corresponds to φ ≈ 299 − 303, or only about four integer values for the threshold. We find numerically that the size of this unstable region decreases when increasing the number of generations *N*, but does not change when varying the aggregation parameter. Dynamical numerical simulations for this range of *θ* are characterized by small oscillations in the number of beetles near the fixed point that can last for thousands of years. We note that Schreiber (2003) finds similar behaviour where previously stable fixed points can become unstable when oscillating populations have an Allee effect, as oscillations below the Allee threshold can lead to extinction. Given the time scale of these oscillations and the small range of tree resilience leading to this behaviour, this apparent bifurcation is unlikely to be important biologically. See Appendix B for more information about the dynamics very near the critical point.

**Fig. 5.**
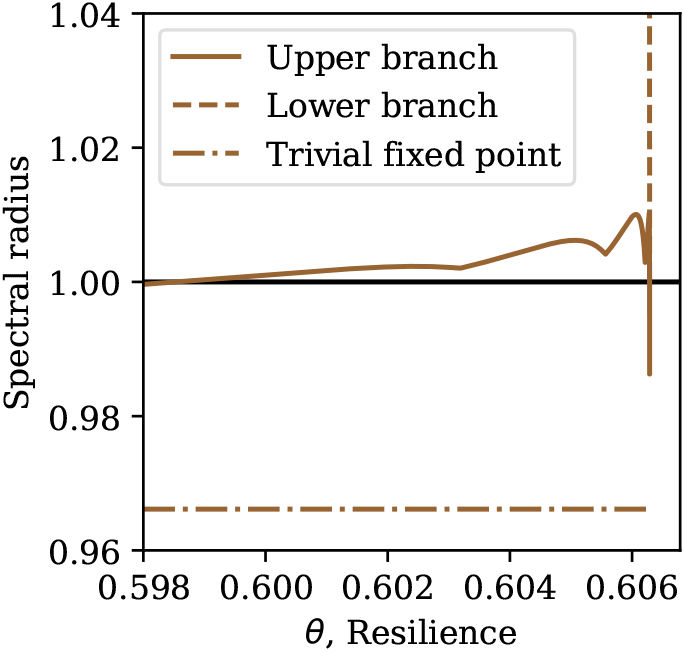
The spectral radius of each fixed point using the Hill approximation and nondimensionalizing to remove *c* as a parameter. We set the aggregation parameter at *κ* = 5

### 5.2 Immigrating beetle population

We now simulate the time dynamics of the model with biologically motivated initial conditions. We initialize a forest stand entirely with juvenile trees and no MPB. This even-aged forest structure could result from large scale forest fires that level entire stands (Safranyik and Wilson 2006). We then introduce MPB into the system at each time step. Biologically, these beetles could represent beetles flying in from an adjacent infested stand, or endemic beetle populations that exist in weakened trees between outbreaks. Eventually, as the trees grow and become susceptible, if there are enough immigrating beetles they cause an outbreak.

We simulate the model for 500 years using the Hill function approximation, though we note that results with the original function are similar. In addition to fixing *N* = 50 and *s* = 0.99 as mentioned above, we fix *κ* = 5, Φ = 200 and *c* = 500 (*θ* = 0.4, though we leave the dimensional parameters). These parameter values ensure that there is a non-trivial stable solution (Fig. 4). We additionally set the total number of trees *T* = 1 so that *j*_*i,t*_ and *S*_*t*_ can be interpreted as fractions of the total number of trees (4). We introduce 50 beetles at each time step to represent beetles that are coming from infested trees outside of the local stand.

We plot the number of susceptible trees and beetles over time in three different ways in Fig 6. We first plot the beetle dynamics over a long time scale in Fig. 6a, then zoom in to the dynamics of a single transient outbreak in Fig. 6b, and finally plot the number of beetles versus the number of trees in Fig. 6c. Note that the number of beetles we plot here does not include the additional immigrating beetles, only the beetles from locally infested trees.

**Fig. 6.**
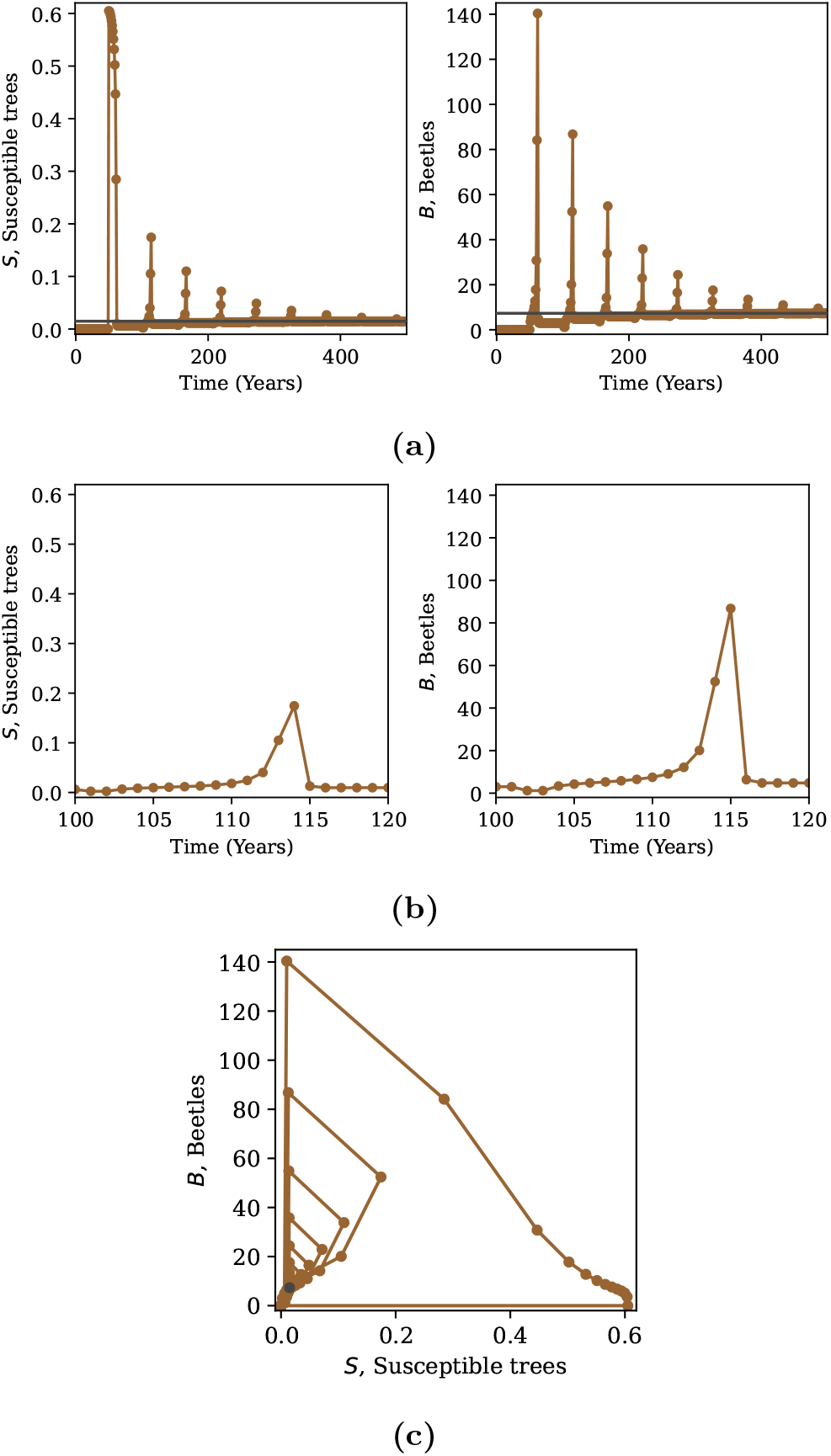
Simulation of the model with the Hill function approximation. We set the parameter values to *κ* = 5, Φ = 200, and *c* = 500 (*θ* = 0.4) with the initial condition *j*_1,0_ = 1 and all other variables set to 0. At each time step, we add 50 beetles that are assumed to migrate from adjacent stands. (a) shows the evolution of the number of susceptible trees *S*_*t*_ and beetles *B*_*t*_ over a very long time with a grey horizontal line indicating the equilibrium values of each, (b) shows the same simulation over a single outbreak, and (c) shows the same simulation again over time in the phase plane with a dark circle overlaid at the equilibrium point. The plotted number of beetles does not include the added immigrating beetles

On a long time scale, Fig. 6a shows that the values of susceptible trees and beetles eventually tend toward their equilibrium values as predicted by the stability analysis (grey lines in Fig. 6a), but that there are large transients in their population sizes. The transients occur because every time there is a peak in the number of susceptible trees from the initial cohort of juveniles, the beetles infest these trees and their population peaks in the following year, reducing the susceptible tree population below its equilibrium value. The beetle population then collapses as they are left with no susceptible trees to infest. We can see this in Fig. 6b.

The transients can be interpreted as outbreaks, and the period is set by the recovery time of the forest. With *N* = 50, the beetle outbreaks peak every 53 years. This is because the susceptible trees peak *N* + 2 = 52 years after they are infested when the seedlings replacing the infested trees become susceptible themselves, and then the beetles peak the following year. Eventually, we reach an equilibrium where the number of successful beetle attacks corresponds exactly to the number of trees that became susceptible that year.

Figure 6c shows another way to visualize the dynamics. Here we start at the origin, with no beetles or susceptible trees, and then the number of susceptible trees increases without changing the number of beetles. The beetles then infest nearly all of the susceptible trees and their population peaks, before collapsing in the following year. This results in a spiral-like pattern as the number of beetles and trees settle to their equilibrium values (marked by a dark circle in Fig. 6c).

The magnitude of the transients can be reduced with alternative initial conditions. If the system is initialized very close to the upper fixed point with a geometric distribution for the juvenile trees, for example, the transient outbreaks are very small as the system settles to equilibrium. However, this requires precise initial conditions that are unlikely biologically, especially given that lodgepole pine is heavily fire dependent.

In addition to beetle outbreak dynamics, our model allows us to analyze the change in forest structure over time. Figure 7 shows the fraction of trees in each age class, including the susceptible trees (marked by *S* on the *x*-axis). We see that the structure changes gradually from the initial condition with all trees as seedlings to the geometric distribution predicted in (36). At each time, there is a peak that represents the cohort of trees regrowing after being killed by MPB. This peak decays over time as the forest reaches its equilibrium structure, which is also when the transient beetle outbreaks end. At very long times, the fraction of trees moving into the susceptible class will be exactly equal to the fraction killed by pine beetle.

**Fig. 7.**
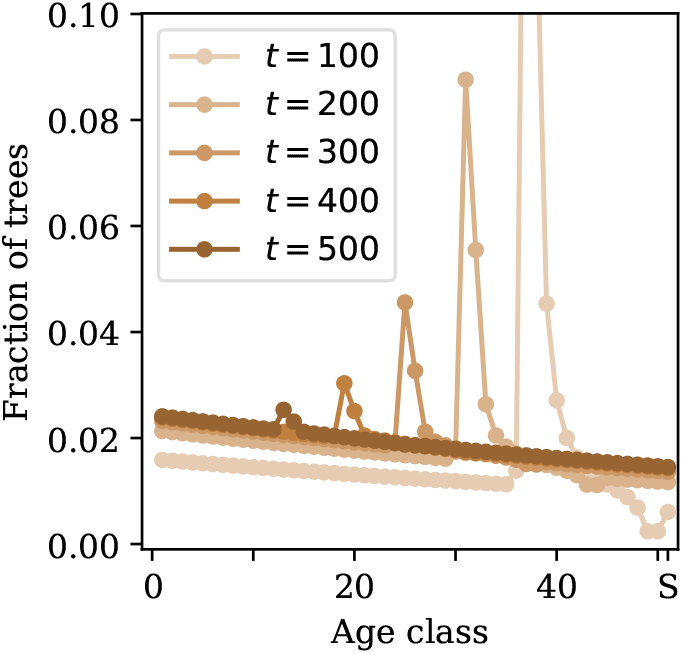
The age structure of the forest over time. The simulation is as in Fig. 6, with the Hill function approximation and parameter values *κ* = 5, Φ = 200, and *c* = 500 (*θ* = 0.4) with initial condition *j*_1,0_ = 1 and all other variables set to 0. At each time step, we add 50 beetles that are assumed to migrate from adjacent stands. The fraction of susceptible trees is shown as the 51st age class

**Fig. 8.**
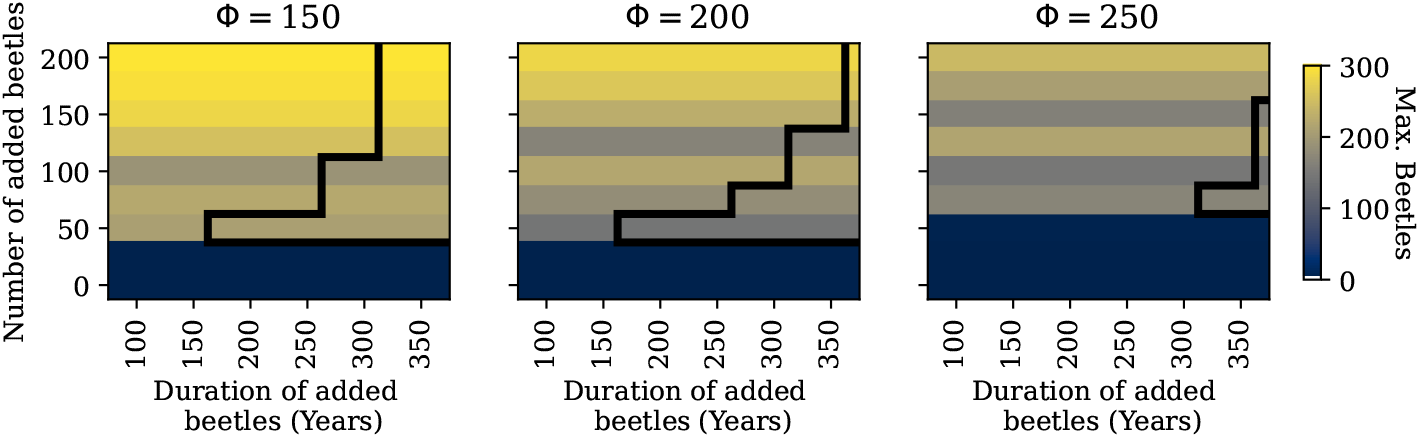
The effects on simulation outcomes of changing the number of beetles immigrating and the duration of their immigration for three values of the thresh-old Φ. We run the simulation for 500 time steps and set it up as in Fig. 6, with the Hill function approximation and parameter values *κ* = 5 and *c* = 500, with initial condition *j*_1,0_ = 1 and all other variables set to 0. We vary the number of beetles added at each time step from 0 to 200 in increments of 25, and we vary the duration of the invasion from 50 to 400 time steps (years) in increments of 25. The colour of the cell indicates the maximum number of beetles present in the stand at any time step (not including the immigrating beetles). The region outlined in black indicates that there is a non-zero beetle population remaining at the end of the simulation takes more transient outbreaks for the population to approach the equilibrium point.

Finally, we consider how these dynamics change as we vary the immigrating beetle population and the tree defense threshold. We run simulations forward in time for 500 years with initial conditions as in the previous simulation in Fig. 6. We again use the Hill function approximation and set *κ* = 5 and *c* = 500, and set the initial condition to *j*_1,0_ = 1 with all other variables set to 0. For each simulation, we set the number of added beetles to be between 0 and 200 in increments of 25. In the previous simulation, we added these beetles at each time step for the entire simulation. Here, we add beetles for a fixed duration that varies between 50 and 400 years in increments of 25. This allows us to test if the added beetles are able to force the system to its non-trivial equilibrium, or if the local beetle population collapses when beetles are no longer added. We do these simulations for three values of the threshold, Φ = 150, 200, and 250.

Figure 8 shows the maximum number of beetles at any point in the run for each combination of parameters. Note that this does not depend on the duration of immigration, which makes sense as the first transient outbreak is the largest (see Fig. 6). It is interesting that the maximum number of beetles does not increase monotonically with the number of immigrating beetles. This is because when the trees first become susceptible there are not enough beetles to consume all of the susceptible trees, and so the beetle population grows for a few time steps (see the first transient in Fig. 6a). At intermediate numbers of added beetles, the beetle population has a few years to grow to a large transient population. At higher number of added beetles, the beetle population peaks the first year after the trees become susceptible as they consume a large enough fraction of the trees that their population decreases after the first year. This makes the first peak in beetle population lower relative to more intermediate numbers of added beetles. When the number of added beetles is very large, then the first transient can be even larger as the beetles consume a very large fraction of the susceptible trees.

Additionally, Fig. 8 outlines a region in black where the beetle population is non-zero at the end of 500 time steps. This effectively means that the local beetle population persists in the absence of added beetles and the system tends towards the upper equilibrium point. In general, the duration of immigration needed for local beetle persistence decreases as the number of added beetles increases. This is because the system settles more quickly to the upper equilibrium when the immigrating beetles cannot immediately consume all available susceptible trees. In other words, as the number of immigrating beetles increases, the susceptible tree population crashes further below the equilibrium value and therefore it

## 6 Discussion

The model we have introduced here combines key elements of MPB population dynamics with a forest age structure and therefore allows us to study the long-term behaviour of MPB. We find that outbreaks in this model are characterized by short transient outbreaks with frequency set by the age of recovery of the forest. In forests where the resilience is above a certain level, MPB populations always die out, but when the resilience is below a threshold level, forests are susceptible to MPB outbreaks. In susceptible stands, decreasing forest resilience decreases the Allee threshold, thereby making them more likely to be infested by MPB.

Insect outbreaks in general are often characterized by large boom and bust cycles (Barbosa and Schultz 1987), and these dynamics are frequently understood mathematically in terms of bifurcation theory where the appearance of new fixed points or the exchange of stability of existing fixed points can lead to sudden changes in population numbers. For example in the models of spruce budworm (Ludwig et al. 1978; Hassell et al. 1999) or experimentally in flour beetles (Cushing et al. 2002). This has also been true in the case of MPB (Heavilin and Powell 2008; Goodsman et al. 2016). In our model, we have shown that there is a fold bifurcation at a critical value of forest resilience where the sudden appearance of additional fixed points can lead to beetle outbreaks (Fig. 4).

In the nondimensionalized version of our model, the forest resilience is described by the parameter *θ*, which is the equal to the threshold for the number of beetles needed to successfully overcome tree defenses divided by the beetle productivity. Thus, a decrease in forest resilience could result either from a decrease in the threshold of beetles needed for a successful attack, which could correspond to weakened trees under drought (Logan et al. 2003; Alfaro et al. 2010), or an increase in the beetle productivity per tree, which could be due to warmer temperatures reducing over winter mortality (Bentz et al. 2022; Safranyik and Wilson 2006). In both cases, climate change is likely to increase the magnitude of these effects. Robbins et al. (2022) recently quantified the effects of climate change with a similar species of bark beetle and found that tree mortality was increased by around 30% because of warming. Understanding how MPB dynamics will continue to shift under climate change will require additional work quantifying its current effects and new modelling efforts to predict these changes in the future.

When there are beetle outbreaks in this model, there are large transients in the number of beetles where the beetle population can briefly reach up to 40 times its equilibrium value (Fig. 8). These transients are larger in magnitude with more uniform tree age distributions as they settle to the geometric distribution at the upper fixed point. The even-aged stand structures that yield larger transient beetle populations are plausible in real forests as large scale fires reset the forest age structure (Safranyik and Wilson 2006), though note that in practice the combination of fire, logging and forestry practices, and MPB leads to complicated forest age structures (Taylor and Carroll 2003; Safranyik and Wilson 2006; Axelson et al. 2009; Axelson et al. 2010). In reality then, it is un-likely that forests will ever settle into the predicted geometric age distribution as it takes hundreds of years for the transient outbreaks to dissipate (Fig. 6a). We note that the recurrent transient outbreaks we see in our dynamical simulations occur because we have introduced an immigrating beetle population. While here we have introduced them continuously for convenience, this is not necessary for recurrent outbreaks. With our initial conditions, after the initial cohort of susceptible trees are infested, there is little regrowth for the next *N* + 2 years. Rather than continuously introducing beetles, we could instead introduce immigrating beetles periodically when there is a larger susceptible tree population. This could represent appropriate conditions for an endemic to epidemic transition, where endemic beetles living in weaker trees are able to take advantage of the new cohort of susceptible trees and then spread to infest new stands (Safranyik and Wilson 2006). Alternatively, we could initialize the forest with a different structure, closer to equilibrium, so that the beetle population is self-sustaining after a short initial introduction. In other words, in a more mixed age forest, a shorter burst of immigrating beetles can lead to recurrent outbreaks, though the outbreaks will be much smaller in magnitude.

The period of the transient beetle outbreaks observed in this model is driven by the age structure of the forest. This type of single generation oscillation, where the period of the oscillation is driven by the time it takes for a cohort of immature individuals to mature and then produce a new cohort of immature individuals, can be seen in many age structured or time delay population models (May 1974; McCauley and Murdoch 1987; Jansen et al. 1990). However, these models typically predict stable oscillations rather than the transient behaviour observed here, which is likely because these models do not include an Allee effect and include overcompensation in addition to a time lag. This means the population crashes at high densities due to overcompensation, but recovers after a time lag as the population can recover even at low densities. Here we require immigrating beetles to overcome the Allee effect, and there is no overcompensation in the beetle dynamics. Despite that, the mechanism is similar, where existing trees suppress new growth until MPB kill enough susceptible trees to clear canopy space for a new cohort of juvenile trees. We note that given the relatively simple juvenile dynamics in this model, it may be possible to translate the model into a lower dimensional system with delays, as Hassell et al. (1999) did with the complicated age structured model of spruce budworm presented in Jones (1979).

The recent outbreaks in Western Canada are largely consistent with the conclusions from our model. In British Columbia, outbreaks have been found to occur every 30 to 40 years (Alfaro et al. 2010; Axelson et al. 2009), which is similar to the outbreak frequency we found with the parameter values in our simulations. We could easily reduce the time between outbreaks by changing the number of years it takes for a seedling to become susceptible, *N* . We note also that Duncan et al. (2015) reduced the time between outbreaks in their model with a variety of methods (including spatial dynamics) that should also be applicable here. In Alberta, MPB populations attacked large areas of novel hosts but have since declined. In the context of this model, this would be the first of a series of transient outbreaks of decreasing amplitude. We would therefore expect large outbreaks to return in less than 50 years as the forests regrow and new host trees become available.

Given that our model is constructed from the models presented in Duncan et al. (2015) and Goodsman et al. (2016), we compare our qualitative results to those models. We first compare with the model in Goodsman et al. (2016), where we note that our model is identical to that model when trees are not replaced, ie. when *s* = 0. The purpose of extending their model was to be able to answer questions on longer time scales, and so here we compare the dynamics of a single outbreak. They find that with very strong aggregation and appropriate initial conditions, the mean number of beetles per tree becomes fixed and then slowly decreases to 0. In our model, the mean number of beetles per tree would also go to the upper fixed point, but it would remain there as the trees infested would be balanced by the new trees entering the susceptible class. Thus, with very strong aggregation, we find that tree replacement is on a relevant time scale as beetles are able to aggregate effectively on even a small number of susceptible trees and the beetle population does not decrease. With weaker beetle aggregation, beetle populations go extinct in very few time steps even if starting above the Allee threshold in the Goodsman et al. (2016) model, and so outbreaks are very short in time. In our model, tree replacement can extend these outbreaks to more biologically reasonable lengths.

The qualitative behaviour of our model is quite different from the behaviour of the Duncan et al. (2015) model. That model does not have an Allee effect, something we know to be important for MPB (Boone et al. 2011). They find cyclic outbreak solutions resulting from overcompensation, where beetles outbreak approximately every 120 years. We do not find any oscillatory solutions in our model, and instead beetle outbreaks are transients with magnitude dependent on the initial conditions and with a frequency set by the age of recovery of the forest. Additionally, the outbreaks in our model are shorter in duration with a faster collapse of the beetle population and tend to produce fewer infested trees than the cyclic ones observed in the Duncan et al. (2015) model.

Finally, we consider several possible extensions of the model introduced here. First, we could extend the model to include a less resilient tree class. MPB maintain endemic populations by attacking trees already weakened by other pests, disease, or age (Raffa and Berryman 1983; Boone et al. 2011; Bleiker et al. 2014). A lower resiliency tree class may allow for beetle persistence between boom and bust cycles without the need for immigrating beetles. Second, a spatial extension of this model would allow us to answer how these transient outbreaks spread over space rather than over time, and would allow us to consider relevant spatial inhomogeneity in pine resilience (Cudmore et al. 2010; Cullingham et al. 2011; Burns et al. 2019). Third, while our survival function *F* captures several key aspects of MPB biology, others could be included. For example, beetles are also able to signal when host defenses have been overcome to prevent further aggregation (Raffa and Berryman 1983; Safranyik and Wilson 2006). This means that the survival function likely overestimates the number of surviving trees with high beetle populations, as when beetles are numerous they do not aggregate as strongly. One option to account for this would be to let *k* be a function of *m*, with higher aggregation at lower beetle number. Using numerical simulations, we find that this makes it easier for beetles to invade at low numbers under certain conditions. Biologically, beetles may also want to avoid too many attacks on a single tree as the beetle larvae have negative density dependence (Raffa and Berryman 1983). We could account for this by modifying the function in (7) to account for this negative density dependence in the same way as in Goodsman et al. (2017), although we would no longer be able to study the system analytically. Finally, on larger spatial scales, beetles may only attack a fraction *α* of trees in the stand, in which case we could assume the beetles are distributed according to a zero-inflated negative binomial distribution in order to derive the function *F* . This leads to a functional response similar to Goodsman et al. (2017) and may allow for a better fit to data depending on the size of the stand.

Long-term models like this one are important given that the focus for MPB management in recent years has moved from intensive management of single outbreaks to long-term forest management and risk assessment. The model we developed here allows us to study how changes in forest resilience will affect long-term forest and MPB dynamics. This will be increasingly relevant as MPB continues to spread to novel hosts and as host resiliency is expected to decrease under climate change.

## Acknowledgments

The authors would like to thank all Lewis Research Group members, Janice Cooke, and both anonymous reviewers for their valuable feedback on this project. Funding for this research has been provided through grants to the TRIA-FoR Project to ML from Genome Canada (Project No. 18202) and the Government of Alberta through Genome Alberta (Grant No. L20TF), with contributions from the University of Alberta and fRI Research (Project No. U22004). This work was supported by Mitacs through the Mitacs Accelerate Program, in partnership with fRI Research. MB acknowledges the support of the Natural Sciences and Engineering Research Council of Canada (NSERC), [PDF – 568176 -2022].

## A Small *k* approximation for *F*

For aggregation levels that are very high, *k* ≪ 1, we can expand the survival function (9) around *k* = 0 to obtain the following approximation

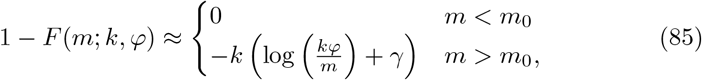

where *γ* is the Euler gamma number, and we can determine *m*_0_ by solving 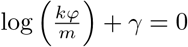to obtain *m*_0_ = *kφe*^γ^ .

The analysis that follows is similar to that using the Hill function approximation, although note that in this case the equation for the fixed points (32) can be solved directly in terms of the Lambert W function. In other words, with this approximation, *m/c* = 1 − *F* (*m*; *k*, φ) can be solved for *m* to obtain

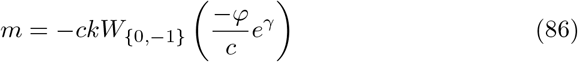

where the *W*_0_ branch is the intermediate (unstable) solution and the *W*_*−*1_ branch is the larger (predominately stable) solution. As *φ/c* → *e*^*−γ−*1^, the two solutions converge to a single unique value 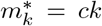 Additionally, for *φ/c* ≤ *e*^*−γ−*1^ ≈ 0.205 there is no solution (because *W* (*x*) does not have real solutions for *x <* −1*/e*). This means that the threshold must be less than about 20% of the number of beetles emerging per tree for the existence of the outbreak equilibrium. The stability analysis of these fixed points follows similarly.

## B Stability near the bifurcation point

Analytically, using the Hill function approximation, we find that there is a fold bifurcation at *m* = *m*^*∗*^ (44), and that the upper branch of the solution very near the bifurcation point must be stable. However, in Fig. 4 the upper branch appears unstable numerically near the critical *m*^*∗*^. Here we show numerically that very near the bifurcation point the upper branch is indeed stable, but that it very quickly loses stability before regaining it at a slightly smaller value of *θ*. The dynamical simulations near equilibrium for values of *θ* in this regime show that the mean number of beetles *m* oscillates rapidly before eventually stabilizing or losing stability. However, even in the case where stability is lost, the simulations often show positive numbers of beetles over times much longer than those relevant biologically.

Figure 9 shows the spectral radius of the upper branch very near the bifurcation point. This is similar to Fig. 5 in the main text, but over a much smaller range of *θ* and only for the upper branch. The second subplot shows that the spectral radius is less than one only very near the critical point, before quickly increasing to greater than one. We additionally plot the real and imaginary parts of the dominant eigenvalue separately in Fig. 10, which are both positive over this range of *θ*. As neither are zero when the spectral radius crosses one, we suspect the system may undergo consecutive Neimark-Sacker bifurcations wherein the oscillations around the fixed points either spiral towards or away from the fixed point.

**Fig. 9.**
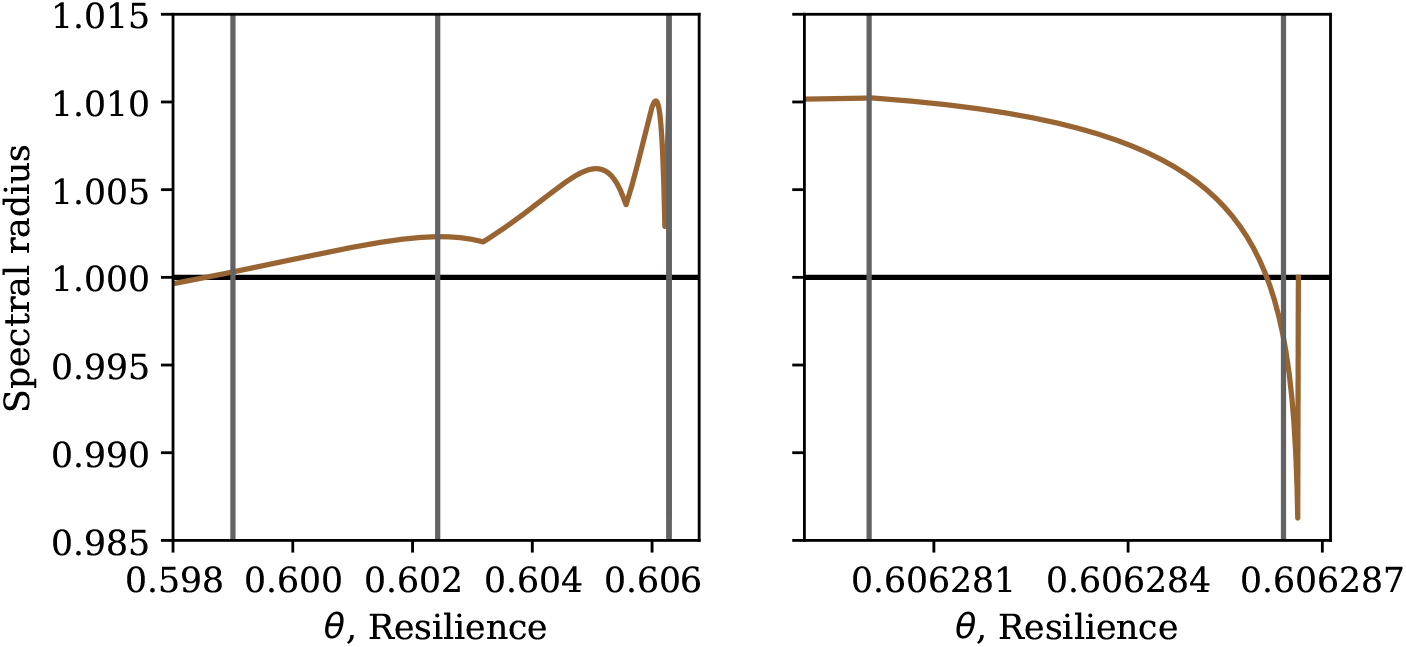
The spectral radius of the upper branch near the bifurcation point with *κ* = 5. The vertical lines indicate the values of *θ* chosen for the dynamical simulation in Fig. 11. The subplots show different ranges of *θ*, where the second subplot is very close to the critical point. Note that the two vertical lines in the second subplot overlap in the first subplot

**Fig. 10.**
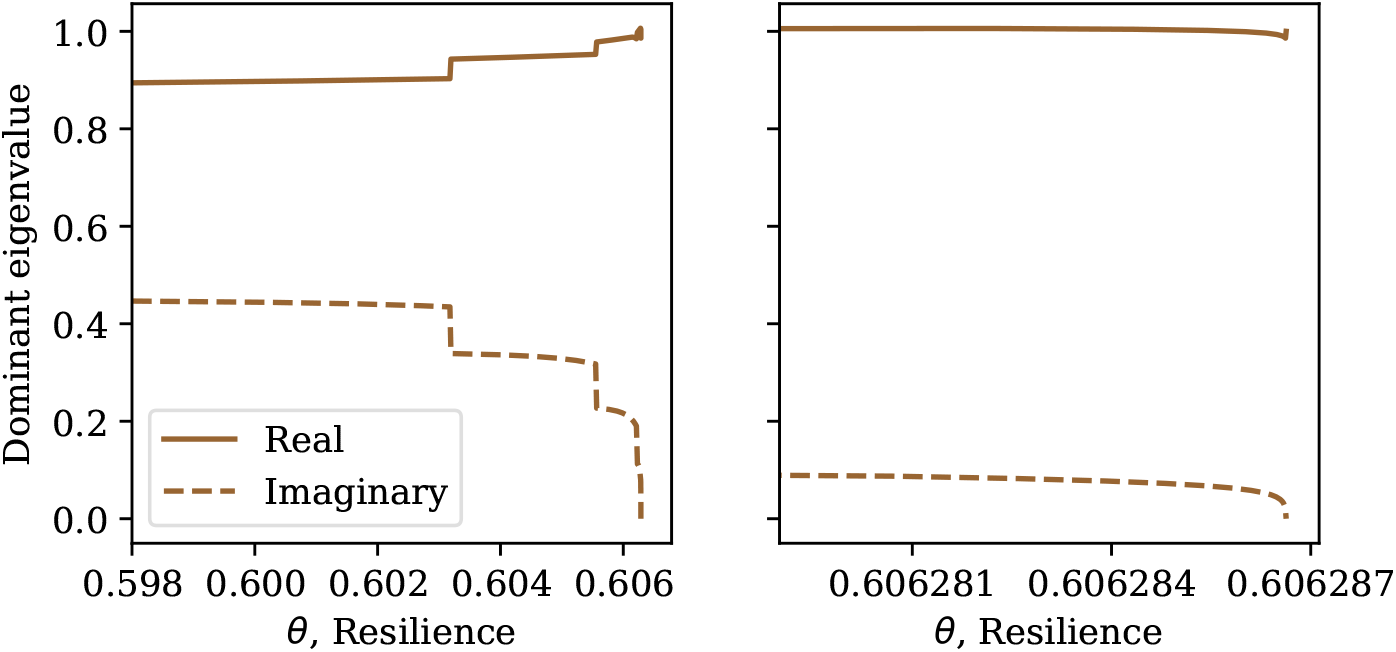
The real and imaginary parts of the dominant eigenvalue of the upper branch near the bifurcation point with *κ* = 5. The subplots show different ranges of *θ*, where the second subplot is very close to the critical point

We simulate the system for four values of *θ* to test what the dynamics of the system look like very near the bifurcation point in Fig. 11. We set the first of these values of *θ* to be near enough to the critical point to be in the stable region (halfway between where the spectral radius crosses one and the critical value of *θ*^*∗*^). We set the next values of *θ* to be at the maximum value of the spectral radius, halfway between the values of *θ* where the spectral radius crosses one, and very near the final point where the spectral radius crosses one. These values of *θ* are shown as vertical lines in Fig. 9. We set the initial values of all parameters and variables to their critical values and simulate forward in time for thousands of years. Note that unlike the simulations in the main text, there are no additional immigrating beetles in this case as we start the simulation close to equilibrium.

**Fig. 11.**
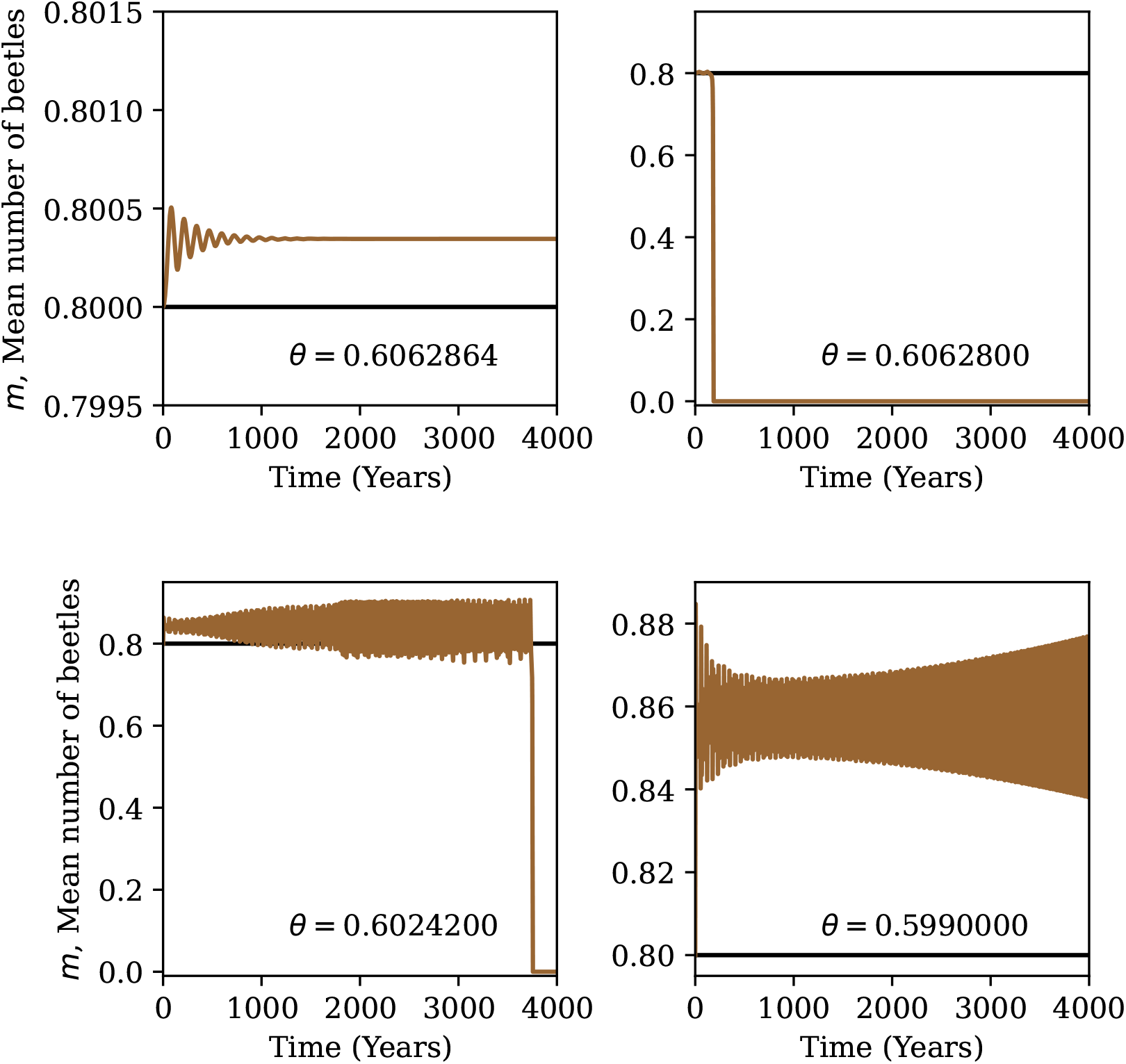
The mean number of beetles from dynamical simulations near the bifurcation point with *κ* = 5. We set the initial conditions to the critical values for all variables. The value of *θ* is given in each subplot and is also indicated in Fig. 9. The horizontal line is the critical value of *m* = *m*^*∗*^ = (*κ* − 1)*/κ*

The first simulation very near the bifurcation point does indeed settle to the stable mean number of beetles in a damped oscillation. For the value of *θ* where the spectral radius is at its maximum, the number of beetles remains non-zero for around 200 time steps (years) before collapsing. For the next two values of *θ*, even though the spectral radius is greater than 1, the mean number of beetles oscillates consistently for thousands of years. Thus, even though this fixed point is technically unstable, it is nearly stable for biologically relevant time scales.

For the second largest value of *θ*, we additionally test how this time to extinction depends on the initial mean number of beetles and plot the results in Fig. 12. We find that small variations in the initial conditions can lead to thousands of time steps difference for the time to extinction. However, as long as there are enough beetles initially, the time to extinction is long enough that the beetle population is effectively stable for biologically relevant time scales.

**Fig. 12.**
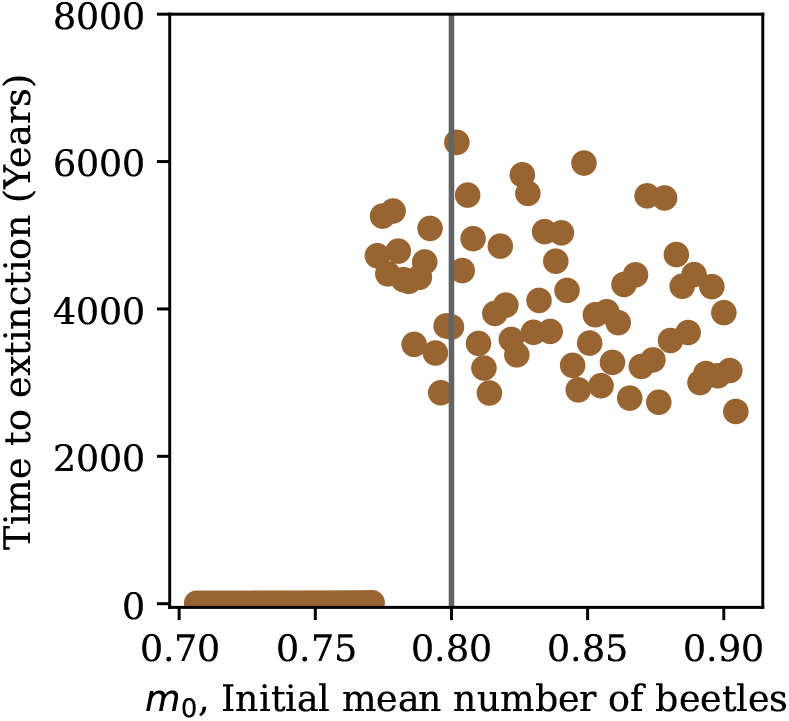
The time to extinction for the beetle population with small variations in the initial mean number of beetles at *θ* = 0.60242. The vertical line shows the critical point *m*^*∗*^. Note that when the initial mean number of beetles is small, the time to extinction is very short, between 6 and 13 years for the range considered here

